# Unraveling the roles of *Mast4* in amelogenesis via regulating DLX3 and stem cell maintenance of mouse incisors

**DOI:** 10.1101/2021.12.15.472878

**Authors:** Dong-Joon Lee, Pyunggang Kim, Hyun-Yi Kim, Jinah Park, Seung-Jun Lee, Haein An, Jin Sun Heo, Min-Jung Lee, Hayato Ohshima, Seiya Mizuno, Satoru Takahashi, Han-Sung Jung, Seong-Jin Kim

## Abstract

Asymmetric division of stem cells allows for maintenance of the cell population and differentiation for harmonious progress. Developing mouse incisors allows for examination of how the stem cell niche employs specific insights into essential phases. Microtubule-associated serine/threonine kinase family member 4 (*Mast4*) knockout (KO) mice showed abnormal incisor development with weak hardness as the apical bud was reduced and preameloblasts were shifted to the apical side, resulting in Amelogenesis Imperfecta. In addition, *Mast4* KO incisors showed abnormal enamel maturation, and stem cell maintenance was inhibited as amelogenesis accelerated. Distal-Less Homeobox 3 (DLX3), known to be a critical factor Tricho-Dento-Osseous (TDO) syndrome, is considered to be responsible for Amelogenesis Imperfecta in humans. MAST4 directly binds to DLX3 and induces phosphorylation at three residues within the nuclear localization sites (NLS) that promote the nuclear translocation of DLX3. MAST4-mediated phosphorylation of DLX3 ultimately controls the transcription of DLX3 target genes, which are carbonic anhydrase and ion transporter genes involved in the pH regulation process during ameloblast maturation. Taken together, our data reveal a novel role of MAST4 as a critical regulator of ameloblast maturation, which controls DLX3 transcriptional activity.

## Introduction

During tooth development, ameloblasts derived from dental epithelial cells and odontoblasts from cranial neural crest mesenchymal cells are responsible for enamel and dentin formation, respectively. Rodent incisors can grow throughout their lifespan and therefore represent a fascinating model for studying the molecular and cellular events involved in stem cell maintenance and differentiation. The renewal of the dental epithelium that produces the enamel matrix and/or induction of dentin formation by mesenchymal cells is performed by stem cells that reside in the apical bud of the incisor (Harada, et al., 2002). The three major stages of amelogenesis are the secretory, transition, and maturation stages. Moreover, ameloblast cell-cell attachment, detachment, and cell movement are regulated so that the characteristic rodent decussating enamel rod pattern can form during the secretory stage of amelogenesis (Shin, et al., 2016; Bartlett and Smith, 2013).

*Wnt* signaling plays an important role in regulating cell proliferation, differentiation, and polarity (Rattanawarawipa, et al., 2016; Logan and Nusse, 2004). The canonical *Wnt* pathway mediates signaling by regulating the intracellular level and subcellular localization of β-catenin (Zhan, et al., 2018). An *in vitro* study of dental pulp cells revealed that *Wnt* signaling is downregulated by Distal-Less Homeobox 3 (DLX3) through regulation of DKK1 expression (Zhan, et al., 2018). However, a recent study reported that *Wnt* signaling could also be upregulated by DLX3 through suppression of DKK4 (Sun, et al., 2019). In addition, in the hair follicle, *Dlx3* is located downstream of *Wnt* (Hwang, et al., 2008). As such, DLX3 and *Wnt* signaling have a complex relationship.

*Dlx3* mutation is known to be responsible for not only Amelogenesis Imperfecta (AI) but also Tricho-dento-osseous (TDO) syndrome, which is an autosomal-dominant human disorder characterized by curly hair at birth, enamel hypoplasia, taurodontism, and thick cortical bone (Zhao, et al., 2016; Hyun and Kim, 2009; Lee, et al., 2008). AI is a genetic disorder characterized by morphological and functional defects in tooth enamel formation. Mutations in several genes are responsible for AI in humans, including *Amelogenin* (*Amelx*), *Ameloblastin (Ambn), Enamelin (Enam), Matrix metalloproteinase-20* (*Mmp20*), *Kallikrein-4* (*Klk4*), *Dlx3, WD Repeat domain-72* (*Wdr72*), *Family with Sequence Similarity 83 Member H* (*Fam83h*), and *Fam20a* (Muto, et al., 2012). Among these, DLX3 is a transcription factor that promotes enamel matrix proteins during amelogenesis (Zhang, et al., 2015). On the other hand, a study using ameloblast-specific conditional knockout (KO) mice of *Dlx3* showed that DLX3 regulation of the target genes related to pH regulation for enamel maturation was more important than that of the genes of enamel matrix protein (Duverger, et al., 2017), suggesting that a complex mechanism exists in the role of DLX3 as a transcription factor during tooth development.

Preameloblasts differentiate into secretory ameloblasts that deposit an extracellular matrix consisting of proteins such as amelogenin, ameloblastin, enamelin, tuftelin, and MMP20 (Kang, et al., 2009; Hu and Simmer, 2007; Stephanopoulos, et al., 2005; MacDougall, et al., 1998), and mineralization is initiated. A shift from matrix deposition to resorption occurs at the transitional and maturation stages, as *Mmp20, Klk4* (Stephanopoulos, et al., 2005), *Amelotin* (Moffatt, et al., 2006; Iwasaki, et al., 2005) and *Odam* (Moffatt, et al., 2008) are predominantly expressed. Ablation of *Mmp20* in mice causes enamel to become thin, brittle, and flake off the underlying dentin (Shin, et al., 2016). KLK4 is a serine protease expressed during enamel maturation, and proteolytic processing of the enamel matrix by KLK4 is critical for normal enamel formation. Two proteases are secreted into the enamel matrix of developing teeth. The early protease was MMP20 and the late protease, KLK4. Mutations in *Mmp20* and *Klk4* both cause autosomal recessive AI (Kang, et al., 2009), a condition featuring soft, porous enamel containing residual protein (Lu, et al., 2008).

Protein kinases that alter the functions of their target proteins by phosphorylating specific serine, threonine, and tyrosine residues play a predominant role in the regulation of many intracellular signal transduction cascades that influence important cellular functions, such as DNA replication, cell growth, proliferation, differentiation, survival, and death (Roskoski, 2015). The microtubule-associated serine/threonine kinase family member 4 (*Mast4*) is a human protein kinase identified in 2006 (Sun, et al., 2006). However, very little research has been undertaken on the developmental role of MAST4 and, to date, most studies have been limited to the brain and mental illness (Landoulsi, et al., 2018; Gongol, et al., 2017; Garland, et al., 2008). A previous study was found that the DNA binding activity of DLX3 was regulated by phosphorylation of serine 138 by Protein Kinase C in keratinocytes and that DLX3 stability was regulated by phosphorylation of serine 10 by Protein Kinase A during osteoblast differentiation (Li, et al., 2014; Park, et al., 2001). However, studies on the phosphorylation of DLX3 in tooth development have not yet been conducted.

In this study, we demonstrated that the incisor teeth of *Mast4* KO mice showed significantly different incisor phenotypes by physicochemical and histological methods. Through RNA sequencing analyses of the separated secretory ameloblasts and apical buds of *Mast4* KO mice, we confirmed an increase in amelogenesis-related genes and a decrease in *Wnt* signaling pathway-related genes. In addition, the regulation of nuclear localization of DLX3 by MAST4 seems to have influenced the physiological condition in the incisor enamel matrix. Here, we report a unique role of MAST4, which expresses the ameloblast layer in postnatal growing mouse incisors. These studies highlight the importance of MAST4 as a key regulator of mouse incisor amelogenesis and stem cell maintenance in the apical bud.

## Results

### Incisor morphology, strength and enamel composition of *Mast4* KO mice show AI phenotype

*Mast4* KO mice were generated using the CRISPR/Cas9 system with a targeted deletion of 71 base pairs on Exon1, which resulted in a premature stop codon by frameshift mutation (Figure 1A). *Mast4* KO mice were grossly similar to WT littermates after birth (data not shown). However, the maxillary and mandibular incisor teeth of *Mast4* KO mice began to bend and showed asymmetrical attrition at postnatal week 6 (Figure 1B, C). At 18 weeks, *Mast4* KO mice were all viable, but they possessed opaque mandibular incisors with chalky surfaces, while WT mandibular incisors were transparent and shiny (Figure 1D, E).

**Figure 1.**
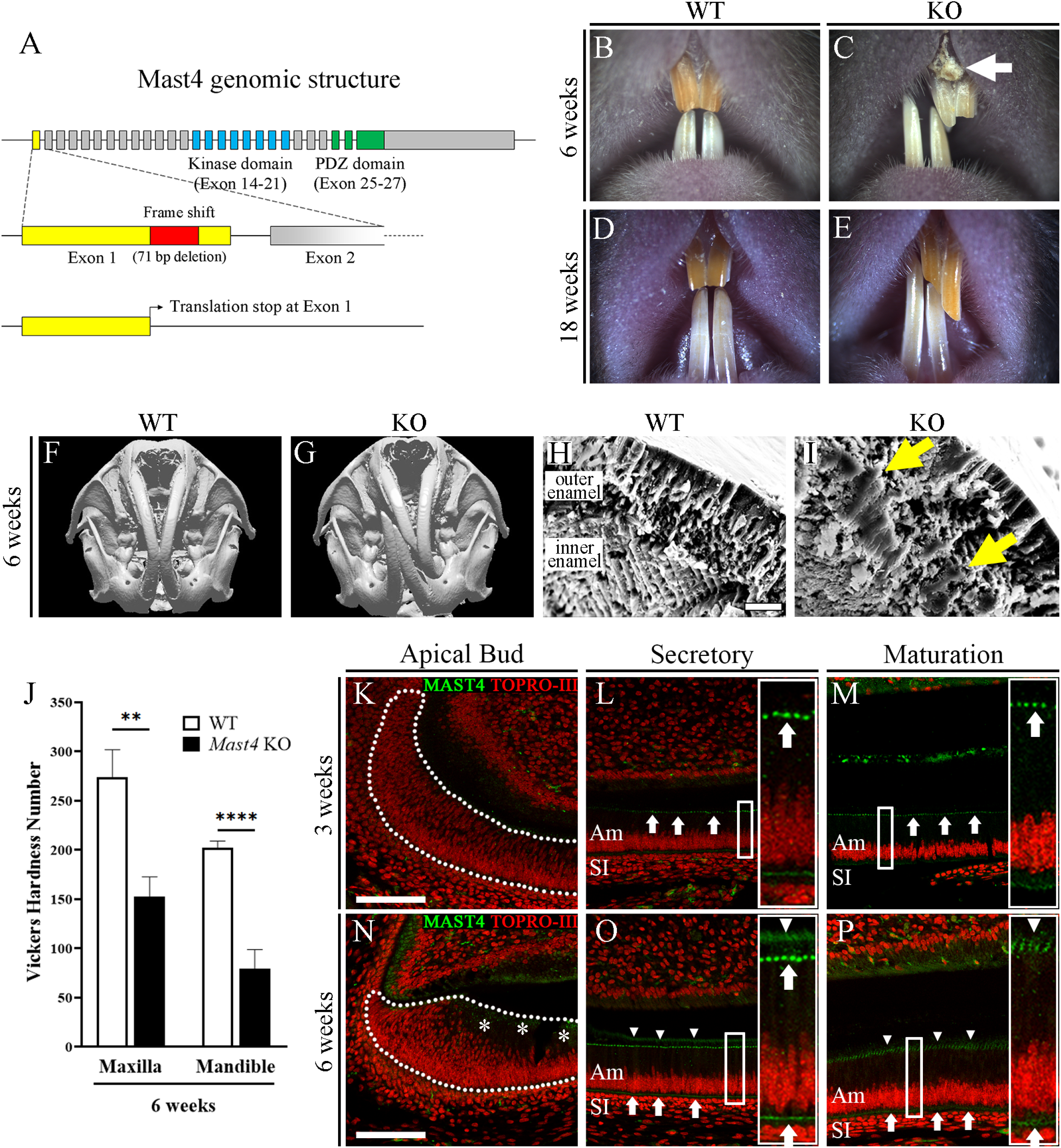
Deletion of *Mast4* leads to amelogenesis imperfecta phenotype. (A) Structure of the *Mast4* KO allele. The mouse *Mast4* locus is depicted with exons as boxes. A new stop codon is generated in the first exon of mRNA after the 71 bp nucleotides are deleted. (B-E) Comparison of 6-week and 18-week WT and *Mast4* KO incisors. Bent maxillary and mandibular incisors are observed in *Mast4* KO mice. An arrow indicates the peeled enamel in 6-week *Mast4* KO maxillary incisor. (F, G) Micro-computed tomography three-dimensional (3D) reconstruction of the *Mast4* KO mouse shows that the incisor is severely curved. (H, I) Scanning electron microscope (SEM) images of incisor enamel dissected planes. Decussated enamel rods are observed in the inner enamel layer of the WT incisor. Enamel rod arrangement is collapsed in the *Mast4* KO incisor (arrows). (J) Vickers microhardness test shows that *Mast4* KO incisors are weaker than WT incisors (N=20 per group, biological replication, detail replication information is described in materials and methods section). Statistical significance was determined with an unpaired *t*-test. *p*-values are: Maxilla=0.0037, Mandible<0.0001. (K-P) MAST4 localization in the WT incisors. The dotted lines indicate the boundary of the apical bud. (K-M) In the 3-week incisor, MAST4 is observed at distal terminal web complexes at secretory and maturation stage ameloblasts (arrows). (N) In 6 weeks, MAST4 expression is extended to the apical side (asterisks). (O) MAST4 is localized at both sides (proximal and distal) of terminal web complexes (arrows) and Tomes’ processes (arrowheads) in the secretory stage ameloblasts. (P) In the maturation stage of ameloblasts, MAST4 is observed at the ruffled border (arrowheads) and proximal terminal web complexes (arrows). Am, ameloblast; SI, stratum intermedium; Scale bars; H, I, 10 μm; K-P, 100 μm.

We also observed twisting to one side between the maxillary and mandibular incisors in 80 to 90% of *Mast4* KO mice, where the incisors were overgrown and the enamel of the maxillary incisors peeled off (Figure 1C arrow). Micro-computed tomography analysis of the head showed incisor malocclusion in *Mast4* KO mice more clearly (Figure 1F, G). To investigate the developmental patterns of molar dentition, histological analysis was performed at the bell stage (embryonic day 18.5; Figure 1 - Figure supplement 1A, B, D, E). The ameloblast layer arrangement of molar tooth germs in *Mast4* KO mice showed an irregular shape (Figure 1 - Figure supplement 1E arrowheads), compared to the molar tooth germs of WT mice. However, there was no difference in morphogenesis, including enamel knot formation. In addition, there were no differences in molar dentition between WT and *Mast4* KO mice until 10 weeks (Figure 1 - Figure supplement 1C, F).

Scanning electron microscopy (SEM) analysis was performed to compare the developing enamel of WT and *Mast4* KO mice (Figure 1H, I). This revealed that the enamel of developing *Mast4* KO mouse incisors contained a collapsed enamel rod arrangement. Furthermore, electron probe microanalyzer (EPMA) analysis was performed to investigate the mineral content of the incisors (Figure 1 - Figure supplement 2A, B). We divided the 6-week incisor calcified portion into two parts (secretory and maturation regions) and proceeded with EPMA analysis. Interestingly, in the maturation part of the *Mast4* KO incisor, calcium (Ca), magnesium (Mg), and phosphorous (P) was reduced, whereas there was no significant difference in the secretory part. We then performed a Vickers test to determine the hardness of WT and *Mast4* KO 6-week incisors. The Vickers test revealed that the loss of MAST4 expression significantly reduced the enamel hardness (Figure 1J). This difference in intensity does not balance the force of the incisor bite and may lead to a twisted phenotype. These results indicated that *Mast4* KO could not facilitate the formation of long and parallel-oriented apatite crystals. Based on these results, the expression pattern of MAST4 (Figure 1K-P) was confirmed in the Tomes process (Figure 1O,P arrowheads) and proximal and distal terminal web complexes (Figure 1L, M, O, P arrows) of the ameloblasts at 3 weeks and 6 weeks in the incisor teeth. At 6 weeks, MAST4 expression was observed at the apical side (Figure 1P asterisks). These results suggest that the weakness of the incisor of *Mast4* KO mice compared to that of WT mice may be caused by dysregulation during amelogenesis.

### Amelogenesis dysregulation in the incisor of *Mast4* KO mice

Apical buds of mouse incisors contain an epithelial stem cell niche that provides the inner dental epithelium (IDE) cells that differentiate into ameloblasts. To investigate stem cell differentiation, the apical bud regions were dissected from the mandibles of 6-week WT and *Mast4* KO mice (Figure2 - Figure supplement 1A, B). Initiation of the enamel was shifted to the apical side of the *Mast4* KO incisor (Figure2 - Figure supplement 1B). Interestingly, the apical bud of the *Mast4* KO incisor appeared to be reduced (dotted line; Figure2 - Figure supplement 1B). In particular, the transit-amplifying (TA) zone seemed to have disappeared. We investigated the histology of the incisor teeth in sagittal, decalcified sections of WT and *Mast4* KO mice at 3 and 6 weeks (Figure 2A-D). No differences in incisor ameloblasts were detected at postnatal day 1 between WT and *Mast4* KO mice (data not shown). In WT mice, secretory stage ameloblasts were tall columnar, and enamel matrix proteins were present in the forming enamel (Figure 2A). During the maturation stage, WT mice exhibited characteristic shortened ameloblasts. In contrast, epithelial cells in the TA zone of the apical bud region appeared to have changed to secretory ameloblasts (Figure 2B). The initiation of enamel matrix secretion in the *Mast4* KO incisor was shifted to the apical bud region, while the initiation of secretion started after the TA region in the WT incisor (Figure 2A, B arrowheads). In the maturation stage, severe hypomineralization was detected in the *Mast4* KO incisor enamel (Figure 2B Maturation). The same features were shown in 6-week WT and *Mast4* KO mice (Figure 2C, D). The enamel layer was detached from the ameloblasts through the apical bud region to the maturation stage region in *Mast4* KO at both stages (Figure 2B, D asterisks). The maturation-stage enamel contained less enamel protein after decalcification in 6-week WT (Figure 2C, maturation arrow). On the other hand, the *Mast4* KO maturation-stage enamel seemed to contain more enamel proteins (Figure 2D Maturation arrow). The enamel matrix proteins are substituted during enamel maturation, and the proportion of proteins in matured enamel is known to be less than 2% m/m(Avery, et al., 2002). This feature of excessive remaining enamel proteins was consistent with the low mineral composition and peeled enamel observed in *Mast4* KO mice (Figure 1C arrow; Figure1 - Figure supplement 2). Ectopic enamel secretion was verified with ameloblastin expression (Figure 2E, F), and ameloblastin was expressed and released to the enamel layer on the apical side in the *Mast4* KO incisor compared to the WT (Figure 2F arrowhead), suggesting that enamel matrix secretion is advanced in *Mast4* KO incisors. Schematic figures present these features of WT and *Mast4* KO incisors and the histological observations are summarized (Figure 2G).

**Figure 2.**
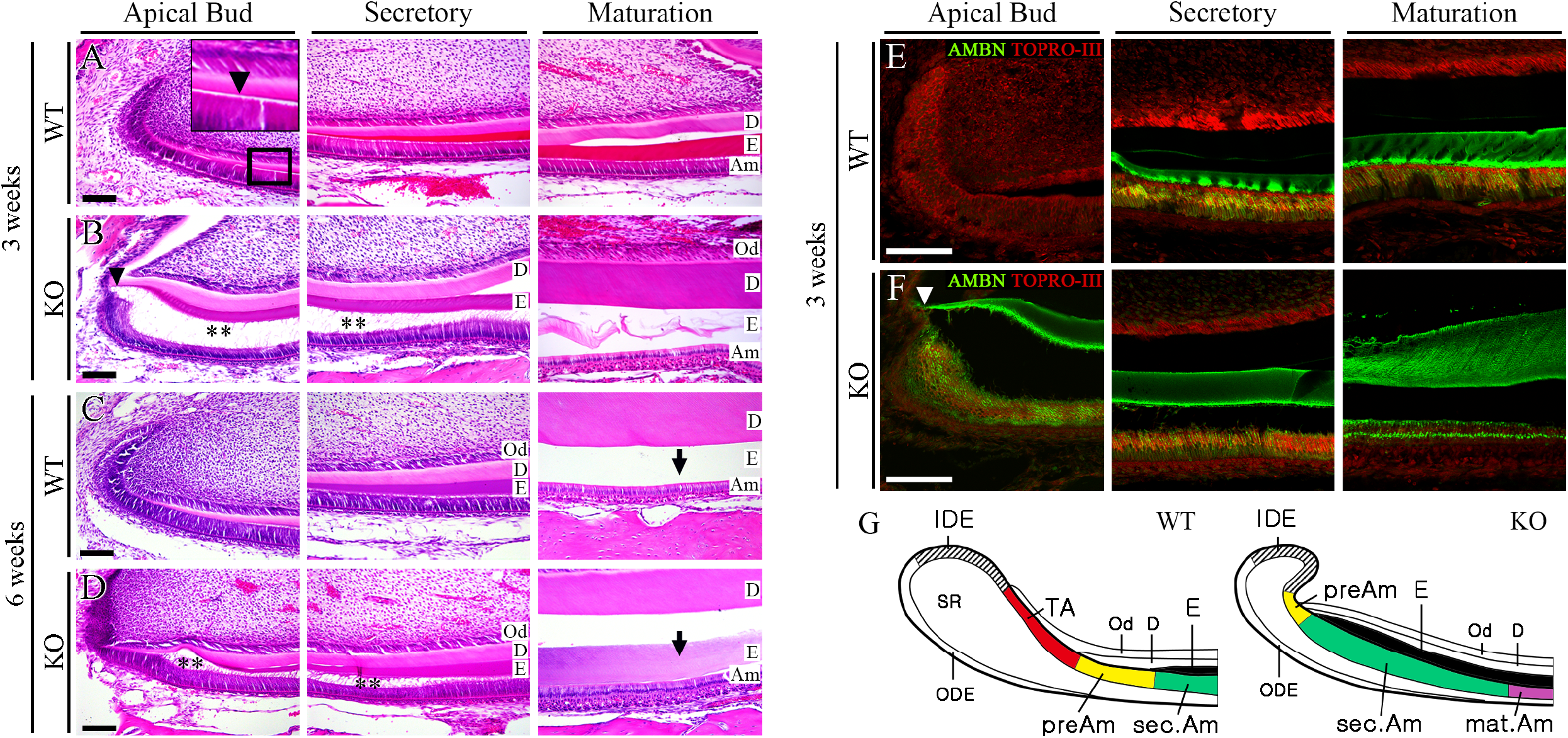
Reduced apical bud region and accelerated ameloblast differentiation in the *Mast4* KO incisor. (A, B) HE stained images in 3-week WT and *Mast4* KO incisors. Arrowheads indicate the initiation position of enamel deposition. (B) In contrast to the WT incisor, detachment is observed in the ameloblasts and enamel of *Mast4* KO incisors (asterisks). The enamel deposition is moved to apical. In the maturation stage, hypomineralization is detected in the *Mast4* KO incisor. (C, D) HE stained images in 6-week WT and *Mast4* KO incisors. (C) After decalcification, the enamel in the WT maturation stage contains less enamel matrix protein than (D) the enamel in the *Mast4* KO maturation stage (arrows). In terms of the amelogenesis stages, early enamel deposition is observed in the *Mast4* KO incisor compared to the WT incisor. The enamel is detached from ameloblasts in the apical bud and secretory stage of the *Mast4* KO incisor (D asterisks). (E, F) Ameloblastin (AMBN) localization in 3-week WT and *Mast4* KO incisor. (F) AMBN is expressed and released to the enamel layer on the apical side in the *Mast4* KO incisor compared to the WT. An arrowhead indicates the initiation position of enamel deposition. (G) Schematic figures of mandibular labial incisor epithelium in WT and *Mast4* KO mouse. In the mandibular incisor of the *Mast4* KO mouse, the labial apical bud including SR and IDE is reduced. TA region is not shown and the initiation of enamel deposition is shifted to the reduced apical bud. D; dentin, E; enamel, Am; ameloblast, Od; odontoblast, sec.Am; secretory ameloblast, mat.Am; maturative Ameloblast, preAm; Preameloblast, IDE; inner dental epithelium, ODE; outer dental epithelium, SR; stellate reticulum, TA; transit-amplifying zone. All scale bars; 100 μm.

To determine and identify additional molecules involved in disrupted amelogenesis in *Mast4* KO mice, we screened MMP20 and FAM83H by immunofluorescence in 6-week incisors derived from WT and *Mast4* KO mice (Figure 3A-H). Interestingly, the expression of MMP20, which aids enamel protein alignment in the enamel matrix and ameloblasts, did not change between WT and *Mast4* KO mice (Figure 3A, B). However, FAM83H secretion to the enamel matrix was significantly increased in the *Mast4* KO secretory region (Figure 3C, D). In the maturation region, the quantity of both proteins contained in the enamel matrix was reduced in *Mast4* KO compared to WT (Figure 3E-H). In particular, the expression of FAM83H, which plays a key role in enamel maturation, shifted to an early stage in the secretory region (Figure 3D, G arrows and arrowheads). These results are consistent with the shifted initiation of enamel secretion and reduced apical bud region in the *Mast4* KO incisor.

**Figure 3.**
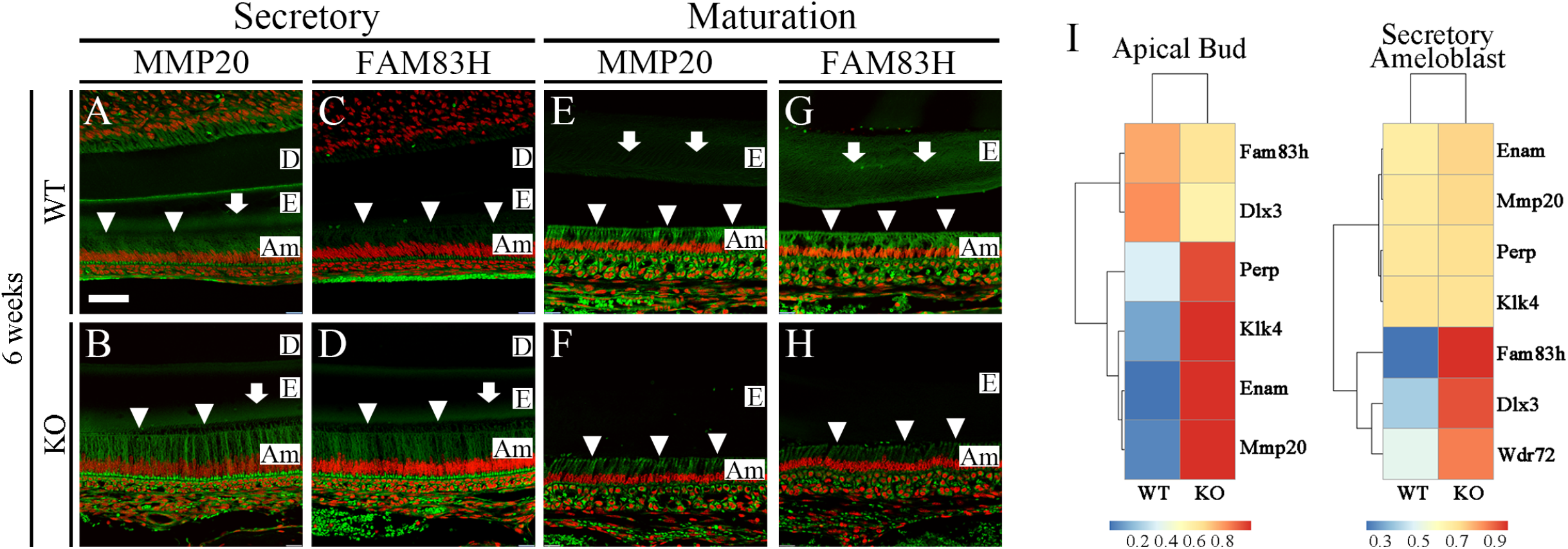
Enamel maturation dysregulation in the *Mast4* KO incisor. (A-H) Enamel matrix protein (MMP20) and maturation-related protein (FAM83H) in the secretory and maturation region of 6-week WT and *Mast4* KO incisors. Arrowheads indicate ameloblasts expressing MMP20 and FAM83H. Arrows indicate the expression of the enamel matrix. (A, B) In the secretory region, the expression of MMP20 is similar between WT and *Mast4* KO. (C, D) FAM83H expression is increased in *Mast4* KO. (E-H) In the maturation region, MMP20 and FAM83H expression is decreased in ameloblasts as well as enamel matrix at *Mast4* KO. (I) RNA-Seq analyses using apical bud and secretory ameloblasts from both the 6-week WT and *Mast4* KO incisor. At apical bud, the expression of *Perp, Klk4, Enam* (*Enamelin*) and *Mmp20* is increased in the *Mast4* KO. At the ameloblast region, the expression of *Fam83h, Dlx3* and *Wdr72* are increased in the *Mast4* KO. D; dentin, E; enamel, Am; ameloblast, Scale bar; 50 μm.

We further investigated RNA sequencing into two regions as apical buds and secretory ameloblasts between 6-week WT and *Mast4* KO incisors (Figure 3I). The major challenge can be considered that the *Mast4* KO apical bud region (properties of secretory ameloblasts and apical bud) and the secretory ameloblast region (properties of maturative ameloblast) due to reduction of apical bud in *Mast4* KO. Consequently, the transcriptome analysis of secretory ameloblasts (Figure 3I) represents exactly matching area as shown in Figure 3A-D. In the apical bud region, enamel matrix protein genes, including *Klk4, Enam*, and *Mmp20*, were increased in the *Mast4* KO mice. In the secretory ameloblast region, enamel matrix maturation-related genes, including *Fam83h, Dlx3*, and *Wdr72*, were increased in the *Mast4* KO mice. These results suggest that enamel secretion and maturation ended at an early stage of amelogenesis in the *Mast4* KO incisor compared to the WT incisor.

### RNA sequencing analysis of *Mast4* deficiency and its effect on the *Wnt* signaling pathway

From RNA sequencing data with separated secretory ameloblasts and apical buds derived from 6-week WT and *Mast4* KO incisors, profile changes in the *Wnt* signaling pathway and stem cell maintenance were found. Genes selected through a differentially expressed gene (DEG) analysis were presented as heatmaps (Figure 4A, C), and a strong decrease in canonical *Wnt* signaling genes and stem cell maintenance-related genes were observed in both apical bud and secretory ameloblast regions. Gene ontology (GO) analysis was performed on the RNA sequencing results (Figure 4B, D). Canonical *Wnt* signaling was significantly reduced in both the apical bud and ameloblasts of *Mast4* KO mice. Interestingly, ossification was downregulated in *Mast4* KO ameloblasts. Some tooth mineralization markers were upregulated in apical buds. We confirmed these changes in the *Wnt* signaling pathway in *Mast4* nulled mHat9d cells, a dental epithelial stem cell line derived from the apical bud epithelium of a mouse incisor, where β-catenin was decreased in both the nucleus and cytoplasm (Figure 4E). *Mast4* ablation markedly decreased the transcription of *Wnt* signaling molecules, including *Wnt-3a, β-catenin*, and *Lrp5* (Figure 4F). This result suggests that ameloblast differentiation is accelerated due to altered *Wnt* signaling and stem cell maintenance in the apical bud region.

**Figure 4.**
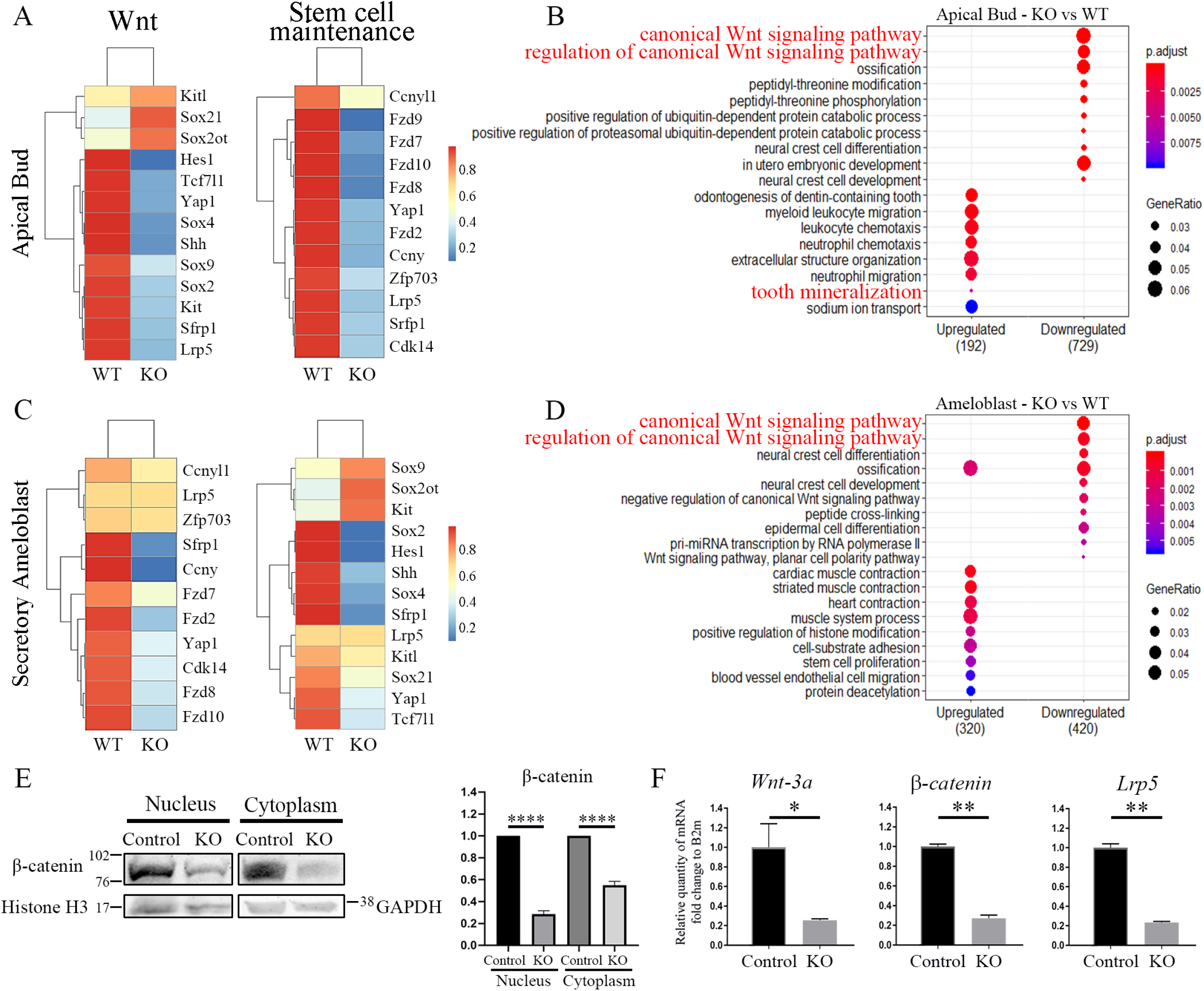
Disruption of stem cell maintenance and *Wnt* signaling pathways in the *Mast4* KO. (A-D) RNA-Seq analyses using apical bud and ameloblasts from both the WT and *Mast4* KO incisor. (A, C) Heatmaps show that canonical *Wnt*-related genes are downregulated in both *Mast4* KO apical bud and ameloblast. Stem cell maintenance related genes are also downregulated in the *Mast4* KO incisor. (B, D) GO analyses at both the apical bud and ameloblast, the canonical *Wnt* signaling pathways are downregulated in the *Mast4* KO incisor. (E) Western blot of β-catenin and its quantification in mHat9d cell line after KO Lentivirus infection (N=4, independent technical replication). Quantification was performed by measuring signal intensities of blot images using ImageJ software and normalized with intensity values of Histone H3 for nucleus and GAPDH for cytoplasm. β-catenin expression in the nucleus and cytoplasm is decreased in the *Mast4* KO cells. Statistical significance was determined with an unpaired *t*-test. *p*-values are: nucleus <0.0001, cytoplasm <0.0001. (F) RT-qPCR analysis of the expression of *Wnt-3a, β-catenin* and *Lrp5* in mHat9d cells (N=3, technical replication). Expression of *Wnt*-related genes is reduced after *Mast4* KO in the mHat9d cells. *p*-values (unpaired *t*-test) are: *Wnt-3a*=0.0483, *β-catenin*=0.0016, *Lrp5*=0.0014.

### MAST4 promoted nuclear translocation of DLX3 by inducing serine/threonine phosphorylation of DLX3 within nuclear localization signal sites (NLS)

Considering that the phenotypes of AI, including enamel disruption and a significant reduction in the mineral contents, shown in *Mast4* KO mice was similar to the phenotypes observed in conditional *Dlx3* KO mice (Duverger, et al., 2017), we focused on examining the relationship between MAST4 and DLX3 transcription factors, which play an essential role in amelogenesis. First, we checked the distribution pattern of DLX3 in WT and *Mast4* KO mice. The expression of DLX3 showed distinct localization patterns depending on the differentiation stages of the incisors in 6-week-old WT mice. DLX3 was predominantly located in the nuclei of ameloblasts at the secretory stage in the incisors of WT mice (Figure 5A arrowheads). At the maturation stage, DLX3 expression still remained high in the nuclei of ameloblasts, although cytosolic DLX3 expression was also increased (Figure 5B, arrowheads). However, in the incisors of *Mast4* KO mice, an even distribution of DLX3 between the cytoplasm and nucleus was observed at the secretory stage (Figure 5C). In particular, DLX3 was predominantly localized in the cytoplasm at the maturation stage (Figure 5D arrows), suggesting that nuclear translocation of DLX3 might be impaired in *Mast4* KO mice.

**Figure 5.**
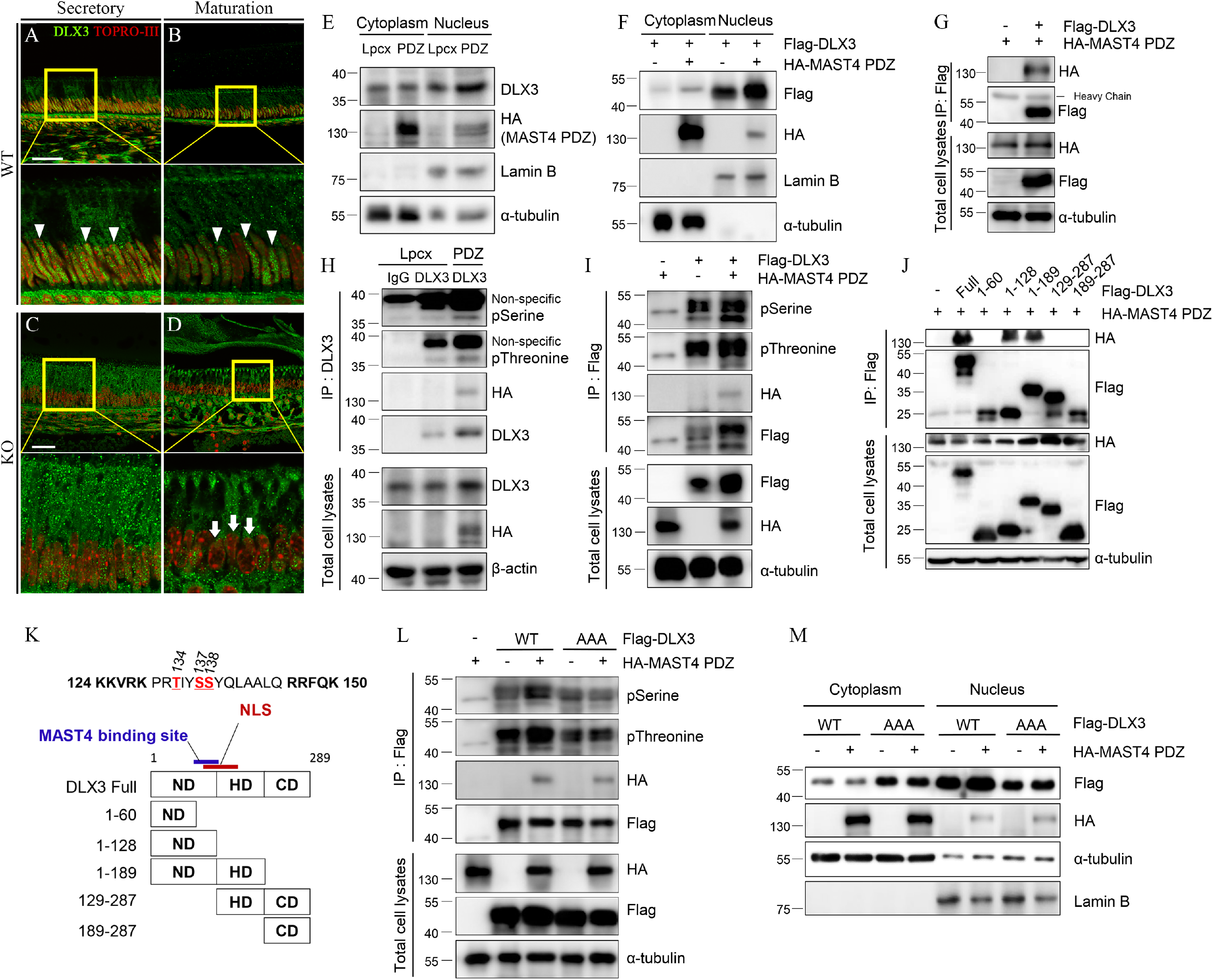
MAST4 regulates translocation of DLX3 by inducing phosphorylation near the NLS region. (A-D) DLX3 localization in 6-week WT and *Mast4* KO incisors. Arrowheads indicate that DLX3 is localized in the nucleus. Arrows point to the nuclei without DLX3. (A) In WT, DLX3 is localized in the nucleus of the secretory stage ameloblast (arrowheads). (B) DLX3 is observed in both the nucleus and the cytoplasm (arrowheads). (C) In *Mast4* KO, DLX3 is distributed in both the nucleus and cytoplasm of ameloblasts in the secretory region. (D) In the maturation stage, DLX3 is localized in the cytoplasm, not in the nucleus (arrows). (E, F) Analysis of DLX3 expression using subcellular fractionation in both PDZ domain overexpressing mHat9d stable cell line and transient transfection in HEK293T. The expression of α-tubulin in the cytoplasm and Lamin B in the nucleus served as controls for the efficiency of subcellular fractionation. (G) Immunoprecipitation assay conducted using HEK293T cells. (H, I) Flag-DLX3 was immunoprecipitated, and the complexes were analyzed by western blot. Note that DLX3 phosphorylation is increased in the presence of MAST4 in mHat9d (H) and HEK293T cells (I). (J) HA-MAST4 PDZ and various deletion mutants of *Dlx3* were co-transfected into HEK293T cells, followed by immunoprecipitation assay. Note that MAST4 binds to the C-terminal of the ND domain. (K) Scheme showing the entire deletion of *Dlx3*. Note that NLS is located in 124-150 aa. (L) Flag-DLX3^WT^, DLX3^AAA^ mutant and HA-MAST4 PDZ were transiently co-transfected into HEK293T cells. Flag-DLX3 was immunoprecipitated, and the complexes were analyzed by western blot. Note that DLX3 phosphorylation is increased in the presence of MAST4. (M) Flag-DLX3^WT^, DLX3^AAA^ mutant and HA-Mast4 PDZ were transiently co-transfected into HEK293T cells and subcellular fractionation was performed. Note that the DLX3^AAA^ mutant has a high proportion in the cytoplasm and is not regulated by MAST4 compared to DLX3^WT^. NLS; nuclear localization signal, ND; N-terminal domain, HD; homeodomain, CD; C-terminal domain, Scale bars; A, B, E, F, 40 μm. **Figure 5 – Source data 1. Uncropped blot images used to generate panels Figure 5E-J**. **Figure 5 – Source data 2. Uncropped blot images used to generate panels Figure 5L, M**.

Based on this observation, we examined whether MAST4 regulated the nuclear translocation of DLX3. Because of its high molecular weight (>285 kDa) and relatively low expression of full-length MAST4, we alternatively used a truncated *Mast4* construct (MAST4-PDZ) that contained a DUF, kinase, and PDZ domain. We first confirmed that DLX3 nuclear translocation was increased when MAST4 was stably overexpressed in mHat9d cells or transiently overexpressed HEK293T cells (Figure 5E and F, respectively). To further explore the relationship between MAST4 and DLX3, an immunoprecipitation assay was performed, and the interaction between MAST4 and DLX3 was observed (Figure 5G). Considering that MAST4 functions as a serine/threonine kinase and that DLX3 phosphorylation affect the DNA binding activity of DLX3 (Park, et al., 2001), we examined whether MAST4 induced DLX3 phosphorylation. Interestingly, MAST4 significantly increased both serine and threonine phosphorylation of DLX3 in both mHat9d and HEK293T cells through immunoprecipitation assay (Figure 5H, I). Next, in order to obtain the basis for MAST4-induced phosphorylation sites, multiple deletion mutants of *Dlx3* were generated, and we found that MAST4 is bound to the C-terminus of the ND domain of DLX3, which was adjacent to the nuclear localization signal (NLS) region (124-150aa) (Figure 5J, K).

In a previous report, it was revealed that the NLS region of DLX3 is a critical location for nuclear translocation of DLX3 (Bryan and Morasso, 2000). In addition, the mechanism by which phosphorylation around the NLS region regulates protein translocation has been elucidated by adopting serine to alanine mutations of the target residues (Chung, et al., 2012; Greco, et al., 2011). Therefore, to investigate whether MAST4-induced phosphorylation adjacent to the NLS of DLX3 is necessary for its nuclear translocation, we generated a non-phosphorylatable mutant with alanine substitutions for the serine or threonine residues (T134, S137, and S138) within the NLS region (DLX3^AAA^). Interestingly, while MAST4 significantly increased serine and threonine phosphorylation of DLX3^WT^, the phosphorylation level of the DLX3^AAA^ mutant was not regulated by MAST4 overexpression (Figure 5L). Furthermore, DLX3^AAA^ was localized in the cytoplasm more abundantly compared to DLX3^WT^, but its nuclear translocation was not promoted by MAST4 (Figure 5M). These results suggest that the NLS phosphorylation of DLX3 by MAST4 is important for the nuclear translocation of DLX3.

### NLS phosphorylation of DLX3 by MAST4 regulated activation of DLX3 target genes involved in pH regulation

To understand the functional implications of NLS phosphorylation of DLX3 transcription factor, a luciferase reporter assay using pGL3-3xDRE, which contained three copies of the DLX3-responsive elements, was performed to assess the transcriptional activity of both DLX3^WT^ and DLX3^AAA^ mutants (Duverger, et al., 2008). As expected, the basal transcriptional activity of DLX3^WT^ was higher than that of the DLX3^AAA^ mutant (Figure 6A). In particular, the transcriptional activity of DLX3^WT^ was further enhanced when MAST4 was overexpressed, whereas the effect of MAST4 overexpression was relatively insignificant to the transcriptional activity of the DLX3^AAA^ mutant. Next, we investigated whether DLX3 occupancy at each target gene promoter was regulated by NLS phosphorylation by MAST4. In a previous report, direct target genes of DLX3, such as carbonic anhydrase and ion transporter genes involved in pH regulation have been identified, and DLX3 binding sites on the promoter of each target gene were identified (Duverger, et al., 2017). Referring to a previous report, ChIP assays were conducted to examine DLX3 binding to the target gene promoter in HEK293T cells. Interestingly, in the case of carbonic anhydrase genes, such as *CA6* and *CA12*, and ion transporter genes, such as *CFTR, SLC24A1*, and *SLC26A1*, DLX3^WT^ showed higher occupancy than DLX3^AAA^ (Figure 6B-F). In particular, the occupancy of DLX3^WT^ on the target gene promoter was further enhanced by MAST4-PDZ overexpression, while that of DLX3^AAA^ was not affected. Next, to confirm whether the mRNA levels of target genes were regulated by altered translocation of DLX3, RT-PCR was performed in mHat9d cells. The mRNA expression of carbonic anhydrases *Car6* and *Car12*, and the ion transporters *Slc26a1* and *Slc34a2* were significantly increased by transfection with DLX3^WT^ and further enhanced when MAST4-PDZ was transiently overexpressed (Figure 6G). However, the mRNA expression of the target genes was not increased by the DLX3^AAA^ mutant, and MAST4-PDZ-mediated increase was not observed when the DLX3^AAA^ mutant was transiently overexpressed. These results indicate that NLS phosphorylation of DLX3 by MAST4 plays an important role in promoting the nuclear translocation of DLX3 and subsequent activation of the target genes.

**Figure 6.**
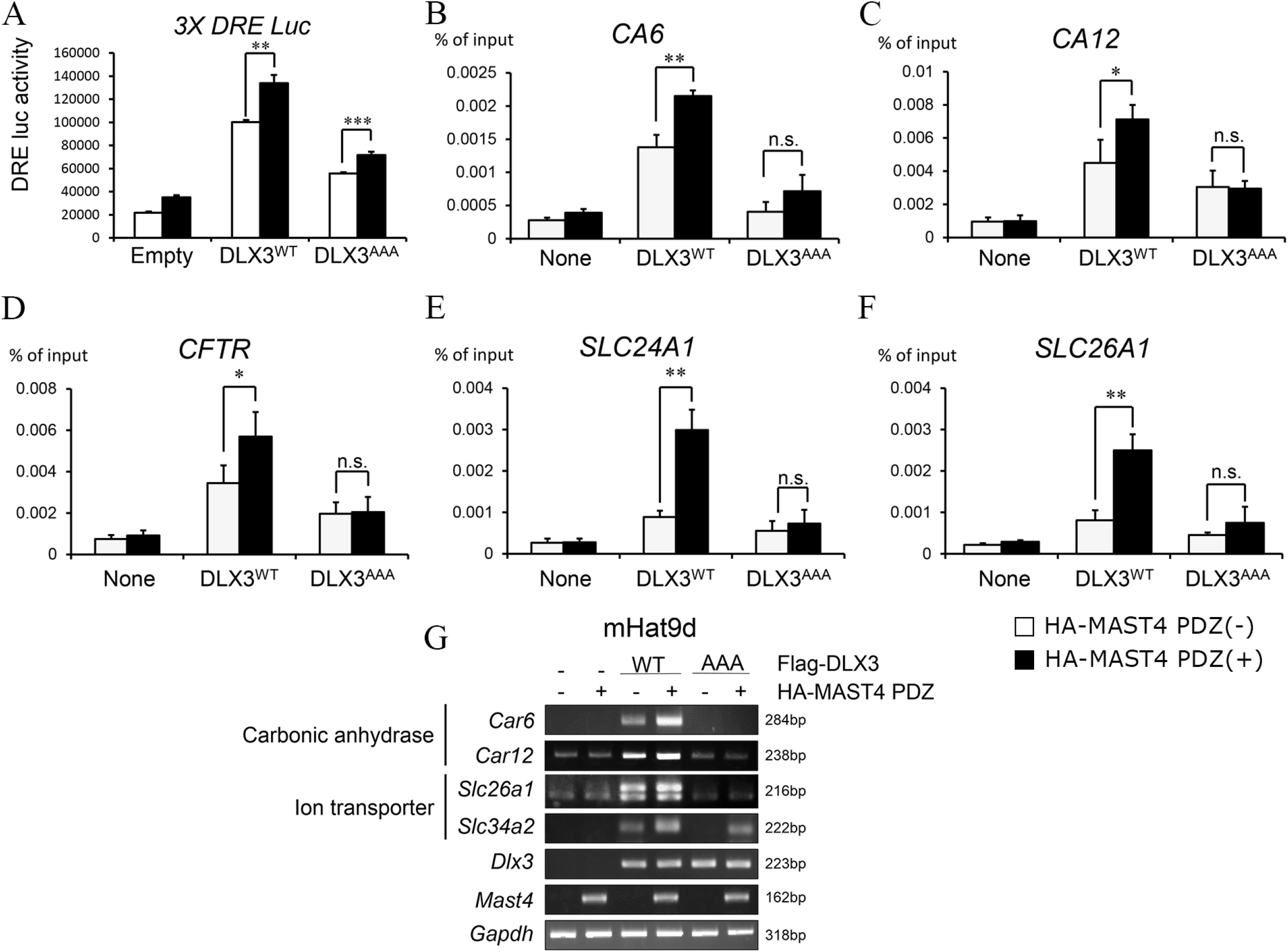
NLS phosphorylation of DLX3 by MAST4 regulates activation of target genes involved in pH regulation. (A) 3x DRE-luc, DLX3, and MAST4 PDZ were transiently overexpressed in the mHat9d cells and beta-galactosidase was co-transfected for normalization. Luciferase activities were measured after 48 h (N=3 per group, technical replication). Statistical significance was determined with an unpaired *t*-test. *p*-values are: DLX3^WT^=0.0013, DLX3^AAA^=0.0009. (B-F) DLX3^WT^, DLX3^AAA^ and HA-MAST4 PDZ were transiently transfected in the HEK293T cells. ChIP assay shows that DLX3^WT^ increased target gene promoter binding and HA-MAST4 PDZ co-transfection further increased whereas DLX3^AAA^ has no significance (N=3 per group, technical replication). *p*-values (unpaired *t*-test) are: *CA6* DLX3^WT^=0.0028, DLX3^AAA^=0.1458; *CA12* DLX3^WT^=0.0454, DLX3^AAA^=0.8724; *CFTR* DLX3^WT^=0.0247, DLX3^AAA^=0.8934; *SLC24A1* DLX3^WT^=0.0021, DLX3^AAA^=0.4967; *SLC26A1* DLX3^WT^=0.0041, DLX3^AAA^ =0.2732. (B-D) Carbonic anhydrases. (E, F) Ion transporters. (G) RT-PCR of carbonic anhydrases and ion transporters involved in pH regulation. **Figure 6 – Source data 1. Measurement data of Luciferase assay of panels Figure 6A**. **Figure 6 – Source data 2. Measurement data of ChIP assay of panels Figure 6B-F**. **Figure 6 – Source data 3. Uncropped gel images used to generate panel Figure 6G**.

## Discussion

*Mast4* KO mice showed abnormal enamel formation in incisor dentition. Specifically, enamel opacity was increased in the mandibular incisors of *Mast4* KO mice, while there was reduced enamel hardness in *Mast4* KO mice compared to WT mice. During dentinogenesis, odontoblasts secrete an unmineralized, collagen-rich extracellular matrix termed predentin. Later, the predentin is transformed into mineralized tissue when apatite crystals are deposited within and around collagen fibrils (Ye, et al., 2004). After dentin matures, enamel can be deposited on the dentin surface by ameloblasts (Nanci, 2017). MAST4 seems to regulate these mechanisms. During amelogenesis, a fibrous protein meshwork and vesicle-like structures were formed instead of the normal enamel matrix in the *Mast4* KO incisor. In several gene transformation studies that produce AI or enamel hypoplasia, abnormal amelogenesis, and disorganization of ameloblasts have been reported (Yan, et al., 2017; Seidel, et al., 2010). The vesicle-like structure formation in the enamel matrix has also been reported in a similar AI induction study (Barron, et al., 2010).

Based on SEM analysis and EPMA testing, the grid shape of the enamel rod array collapsed, and insufficient mineral content was observed. The hardness study also showed that the overall enamel quality was inferior, and this problem is thought to be caused by a failure in the maintenance mechanism of the stem cell niche, which is known to exist in the apical bud. This finding is supported by the fact that this abnormality was not found at the embryo or newborn stage, and no disruption was found in relation to the molar dentition, which does not require stem cell maintenance. Failure of appropriate stem cell maintenance may shift or advance the differentiation of inner dental epithelium cells to a secretory ameloblast stage in the apical bud region. The enamel matrix secretion was also accelerated in sequence, and the discrepancy with maturation timing was thought to cause physicochemical problems.

The present study suggests that DLX3 is a key factor associated with stem cell maintenance and maturation regulation. Several previous studies have shown that DLX3 regulates enamel matrix protein secretion by functioning as a matrix protein transcription factor (Duverger, et al., 2017; Zhang, et al., 2015) and by regulating DKK1, which is known as a *Wnt* signaling inactivator (Yang, et al., 2018; Zhan, et al., 2018). In addition, it has been reported that DLX3 is a downstream target of the *Wnt* signaling pathway in hair follicle development (Hwang, et al., 2008). Interestingly, RNA sequencing data revealed that *Mast4* regulated not only *Dlx3*-related genes, but also canonical *Wnt* signaling-related genes. Numerous studies have shown that *Wnt*/β-catenin signaling is closely related to stem cell self-renewal and maintenance in several organs (Kretzschmar and Clevers, 2017; Xu, et al., 2016; Ring, et al., 2014; Nusse, 2008). Moreover, our previous study revealed a failure of spermatogonial stem cell maintenance in the *Mast4* KO testis, although the signaling pathway involved is different from DLX3 (Lee, et al., 2020). In addition, considering that no abnormalities were found in the molars where adult stem cells do not exist after their development being completed at the fetal stage, it can be inferred that MAST4 is involved in the maintenance of adult stem cells in several signaling pathways.

At the amelogenesis stage, it is well known that DLX3 also regulates transcription of enamel matrix genes. However, it was reported that the transcription of DLX3 target genes related to enamel matrix was not significant in the study using *Dlx3* conditional KO mice, although they still showed the AI phenotype (Duverger, et al., 2017). In our study, the mRNA expression levels of enamel matrix proteins did not change significantly, suggesting that another Dlx3 targets such as genes involved in pH regulation during ameloblast maturation are critical for AI phenotype. Our results demonstrate that MAST4 interacts with DLX3 and regulates DLX3 transcriptional activity, which may ultimately affect incisor amelogenesis. In addition, the nuclear location of various proteins that have NLS is regulated by the phosphorylation of NLS (Nardozzi, et al., 2010). We confirmed that MAST4 is directly bound to DLX3 and phosphorylated three residues located in the NLS, ultimately enhancing both nuclear translocation and target gene activation. The result that the genes involved in pH regulation are specifically regulated by MAST4 and DLX3 is highly correlated with the abnormal ion distribution shown in the previous EPMA results in *Mast4* KO mice (Figure 1 - Figure supplement 2). Overall, the mislocalization of DLX3 due to the loss of MAST4 failed the ameloblast maturation process, which requires appropriate pH regulation, ultimately resulting in the AI phenotype. These previous studies and our results suggest that MAST4 is a putative candidate involved in controlling the activity of DLX3.

Considering that DLX3 is a transcription factor, the distribution of DLX3 appears to affect enamel matrix protein secretion by moving to the nucleus during the secretory stage. The fact that MAST4 is involved in the nuclear localization of DLX3 also supports this hypothesis. A summary of the relationship between MAST4 and DLX3 is shown in Figure 7. Despite the nuclear localization of DLX3 in mHat9d cells, the cause of decreased FAM83H and ameloblastin was not uncovered in this study. Since mHat9d, a cell line originating from the apical bud was used, it suggests the possibility of DLX3 acting differently, depending on the degree of differentiation. Specifically, transcriptome analysis of the previous study reported that the genes involved in pH regulation (*Cftr, Slc24a4, Slc26a7, Slc34a2*, and *Slc39a2*) increased significantly from the secretory stage to the maturation stage, and the expression of *Mast4* also increased by 3.2-fold (Simmer, et al., 2014), raising the need for a more detailed study of *Mast4* during ameloblast maturation.

**Figure 7.**
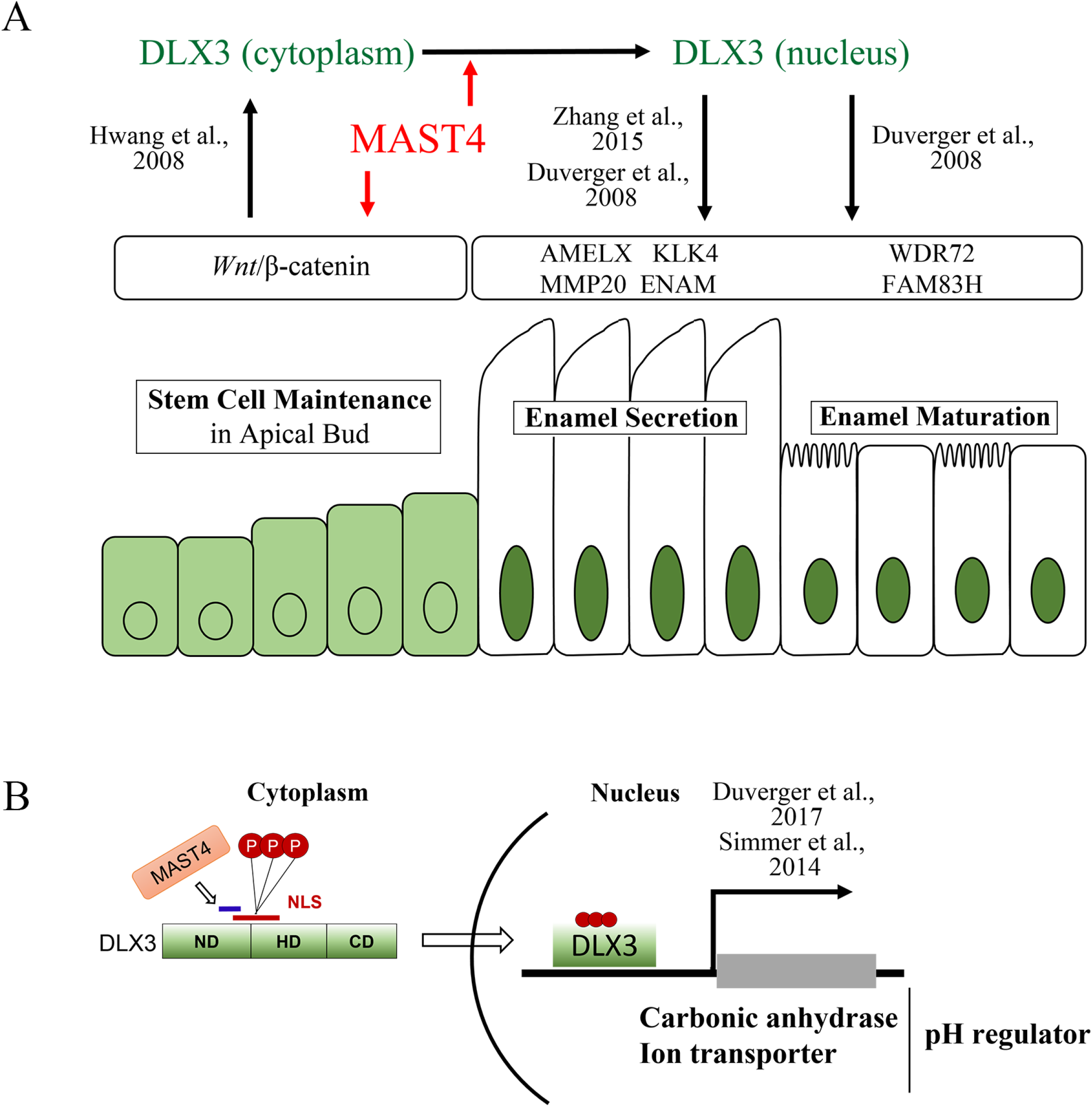
MAST4 regulates amelogenesis in different ways spatiotemporally. (A) MAST4 acts as a stem cell maintenance mediator in incisor apical bud. Whang et al. reported that DLX3 is a downstream target of the *Wnt* signaling pathway. Our RNA sequencing data show that the canonical *Wnt* signaling-related genes are down-regulated in *Mast4* KO incisor tissues. MAST4 functions as a kinase of DLX3 in ameloblasts during enamel maturation via controlling DLX3 nuclear localization. It is well known that *Dlx3* is an Amelogenesis Imperfecta (AI)-related gene. Zhang et al. and Duverger et al. reported that DLX3 regulates the secretion of enamel matrix proteins and enamel maturation proteins by functioning as a transcription factor of those matrix proteins. RNA sequencing data show that *Mast4* regulated *Dlx3*-related enamel matrix and maturation genes. (B) DLX3 controls the expression of pH regulators, one of the critical factors of enamel maturation. Simmer et al. reported DLX3 regulates the expression of carbonic anhydrases and ion transporters which are essential for enamel maturation. Luciferase assay and ChIP data show that carbonic anhydrases and ion transporters are downregulated in *Mast4* KO. Spatiotemporally proper regulation of *Wnt* and DLX3 localization by MAST4 is a key mechanism of stem cell maintenance, differentiation and acquiring physical properties of ameloblast products.

In conclusion, MAST4 plays a key role in the maintenance of stem cells and the regulation of differentiation. The ablation of *Mast4* causes accelerated amelogenesis in the incisor tooth, improper enamel maturation, and abnormal physical properties. These phenomena are triggered by the regulation of DLX3 localization by MAST4. These findings suggest a novel mechanism for controlling the transcriptional activity of DLX3. MAST4 is closely associated with the entire amelogenesis process in mouse incisors and could therefore represent a crucial modulator of AI.

## Materials and Methods

### Animals

All animal experiments were approved by Yonsei University Health System Institutional Animal Care and Use Committee (YUHS-IACUC) in accordance with the Guide for the care and use of laboratory animal (National Research Council, USA). The animal study plan for these experiments (2017-0206) was reviewed and approved by this committee. All the mice were housed in a temperature-controlled room (22°C) under artificial illumination (lights on from 05:00 to 17:00) and 55% relative humidity, and they had ad libitum access to food and water. Adults from each developmental stage (3 weeks, 6 weeks, 10 weeks, and 18 weeks) were used in this study.

To generate *Mast4* KO mice by CRISPR/Cas9-mediated gene targeting, we targeted exon 1 of *Mast4* (RefSeq Accession number: 175171); 5’-GGAAACTCTGTCGGAG GAAG-3’ (exon 1). We then inserted each sequence into the pX330 plasmid, which carried both guide RNA and Cas9 expression units, from Dr. Feng Zhang (Addgene plasmid 42230) (Cong, et al., 2013). We named these vectors as pX330-Mast4-E1 and pX330-Mast4-E15. The schematic figure of the targeted exon1 and translated peptide is illustrated in Figure 1A.

Pregnant mare serum gonadotropin (five units) and human chorionic gonadotropin (five units) were intraperitoneally injected into female C57BL/6J mice (Charles River Laboratories, Kanagawa, Japan) at a 48-h interval, which were then mated with male C57BL/6J mice. The pX330-Mast4-E1 and pX330-Mast4-E15 (circular, 5 ng/μl each) were co-microinjected into 231 zygotes collected from oviducts of mated female mice. The surviving 225 injected zygotes were transferred into the oviducts of pseudopregnant ICR females, and 47 newborns were obtained. Genomic DNA was collected from the tails of 31 surviving founder mice.

To confirm the indel mutation induced by CRISPR/Cas9, we amplified the genomic region, including the target sites, by PCR with the primers for exon 1 target (Supplementary Table 1). The PCR products were sequenced using the BigDye Terminator v3.1 Cycle Sequencing Kit (Thermo Fisher Scientific) and the Mast4-1 genotype F primer. In male founder #38, we found indel mutations in both exon 1 and exon 15 without pX330 random integration. To identify the indel sequence and whether indel mutations in exon 1 occurred on the same chromosome (cis manner), founder #38 was mated with a wild-type female, and the indel mutations in F1 were sequenced. We obtained 17 F1 newborns, of which 12 carried a 71 bp deletion (chr13:103,333,981-103,334,051: GRCm38/mm10) in exon 1 in a cis manner.

### Vickers hardness test

Erupted portions of incisors from 6-week-old WT and *Mast4* KO littermate mice were washed and dehydrated with graded alcohol. Incisors were sagittally embedded in a hard-formulation epoxy embedding medium (EpoFix, EMS, Hatfield, PA, USA). Samples were ground and polished with an EcoMet 30 grinder polisher (Buehler, IL, USA), 1500 grit sandpaper, 400 rpm, and 0.25 μm. The enamel microhardness of the polished samples was measured using an MMT-X testing machine (Matsuzawa, Akita, Japan). Testing was performed with a load of 25 g for 5 s using a Vickers tip. Five indentations per sample were performed on eight incisors (four maxillary and four mandibular) per group and measured.

### Cell culture

The mHat9d cell line was obtained from Professor Harada’s laboratory (Iwate Medical University, Japan). mHat9d is a dental epithelial stem cell line derived from the apical bud epithelium of a mouse incisor. Cells were cultured in 1:1 mixture of Dulbecco’s modified Eagle’s medium and Ham’s F-12 (DMEM/F-12; #11320-033, Life Technologies, USA) containing B-27 supplement (#17504-044, Life Technologies, USA), Fibroblast Growth Factor-basic (bFGF; 25 ng/mL, #100-18B, PeproTech, Inc., USA), Epidermal Growth Factor (EGF; 100 ng/mL, AF-100-15, PeproTech, Inc., USA) at 37°C in a humidified atmosphere with 5% carbon dioxide (CO2). The HEK293T cells were grown in DMEM (WELGENE, Korea) containing 10% fetal bovine serum (WELGENE) and 1% penicillin-streptomycin (WELGENE).

To establish *Mast4*-depelted mHat9d cells, lentiCRISPRv2 vector (#52961, Addgene, USA) was digested with BsmBI and ligated with annealed oligonucleotide targeting *Mast4* exon 1, 5’-TACCCTGCCGCTGCCGCACC-3’ (LentiCRISPRv2-Mast4 Ex1). The vector without insertion was used as a control. To generate the lentivirus, HEK293T cells were transfected with LentiCRISPRv2-Mast4 Ex1 and packaging vectors (pVSVG and psPAX2) using FuGENE® (E2311, Promega, WI, USA) at 70% confluency. The viral supernatant was harvested at 48 h post-transfection, filtered through 0.45-μm filters and applied to mHat9d cells. The clonal cells were selected with puromycin (A11138-03, Life Technologies, USA) at 48 h post-transfection. To establish retrovirus based-Mast4-overexpressing mHat9d cells, both control LPCX and MAST4-PDZ-LPCX vectors with pVSVG were transfected in GP2-293 cells. The supernatant containing recombinant retroviruses was collected 36 hours after transfection and filtered through 0.45-μm sterilization. Virus application and puromycin selection were performed using the same protocol as for lentivirus.

### RNA sequencing

Libraries were prepared for 150 bp paired-end sequences using a TruSeq Stranded mRNA Sample Preparation Kit (Illumina, CA, USA). Namely, mRNA molecules were purified and fragmented from 1 μg of total RNA using oligo dT magnetic beads. The fragmented mRNAs were synthesized as single-stranded cDNAs through random hexamer priming. By applying this as a template for second strand synthesis, double-stranded cDNA was prepared. After the sequential process of end repair, A-tailing and adapter ligation, cDNA libraries were amplified with PCR (Polymerase Chain Reaction). The quality of these cDNA libraries was evaluated with the Agilent 2100 BioAnalyzer (Agilent, CA, USA), and they were quantified with the KAPA library quantification kit (Kapa Biosystems, MA, USA) according to the manufacturer’s library quantification protocol. Following cluster amplification of denatured templates, sequencing was progressed as paired-end (2×150bp) using Illumina NovaSeq 6000 sequencer (Illumina, CA, USA). Low quality reads were filtered according to the following criteria: reads contain more than 10% skipped bases, reads contain more than 40% of bases whose quality scores are less than 20 and reads with an average quality score of less than 20. The whole filtering process was performed using the in-house scripts. Filtered reads were mapped to the reference genome related to the species using the aligner TopHat (Trapnell, et al., 2009). The gene expression level was measured with Cufflinks v2.1.1 (Trapnell, et al., 2010) using the gene annotation database of the species. To improve the accuracy of the measurement, multi-read-correction and frag-bias-correct options were applied. All other options were set to default values.

### Subcellular fractionation and Western blot

Virus-infected cells were lysed in RIPA buffer containing a protease inhibitor cocktail (cOmplete; #11697498001, Roche, IN, USA). Nuclear and cytoplasmic fractions of mHat9d cells were performed using the NE-PER Nuclear and Cytoplasmic Extraction reagents (Thermo Scientific), according to the manufacturer’s protocols. Cell extracts were fractionated by SDS-PAGE transferred to a polyvinylidene difluoride membrane using a transfer apparatus according to the manufacturer’s protocols (Bio-Rad). After incubation with 3% BSA in TBST (10 mM Tris, pH 7.4, 150 mM NaCl, 0.1% Tween 20) for 60 min, the membrane was incubated with antibodies against anti-β-catenin (SC-7199, Santa Cruz Biotechnology, Inc., USA; dilution 1:2000), anti-Histone H3 (ab4729, Abcam, UK; dilution 1:200) and anti-GAPDH (SC-32233, Santa Cruz Biotechnology, Inc., USA; dilution 1:100), anti-Flag (F3165; Sigma-Aldrich, USA; 1:5000), anti-HA (sc-7392, Santa cruz Biotechnology, USA; 1:1000), anti-Lamin B (SC-374015, Santa Cruz Biotechnology, Inc., USA; 1:1000) and anti-α-tubulin (T5168, Sigma-Aldrich; 1:3000) at 4 °C overnight. Membranes were washed six times for 10 min and incubated with HRP-conjugated secondary antibodies for 2h. Blots were washed six times with TBST and developed with the ECL system (RPN2232, GE Healthcare Life Sciences, USA) according to the manufacturer’s protocols.

### Immunohistochemistry

Samples were fixed in 4% paraformaldehyde in phosphate buffered saline (PBS), decalcified in 10% EDTA (pH 7.4; BE021, Bio-solution co. Ltd., Korea) for 48 h at 50°C and then embedded in paraffin using standard procedures. Sections (6-μm thickness) of the specimens were incubated in 10 mM citrate buffer (pH 6.0) overnight at 60°C or Proteinase K (10 μg/ml, AM2546, Thermo Fisher Scientific, United States) for 20 min at 37 °C. The specimens were incubated with anti-MAST4 (BS5791, Bioworld Technology, Inc., USA; dilution 1:150), anti-DLX3 (PA5-40506, Invitrogen, OR, USA; dilution 1:100), anti-MMP20 (ab198815, Abcam, UK; dilution 1:100), anti-FAM83H (NBP1-93737, Novus Biologicals, USA; dilution 1:50) antibodies at 4°C overnight. The specimens were incubated with goat anti-rabbit Alexa Fluor 488 (A11008, Invitrogen, OR, USA; dilution 1:200) or goat anti-mouse Alexa Fluor 488 (A11001, Invitrogen, OR, USA; dilution 1:200). The sections were counterstained with TO-PRO™-3 (T3605, Invitrogen, OR, USA; dilution 1:1000) and examined using a confocal laser microscope (DMi8, Leica, Germany).

### Immunoprecipitation (IP)

Cells were lysed with RIPA buffer containing protease inhibitor cocktail (Complete; Roche). For immunoprecipitation, protein extracts were incubated overnight with the indicated primary antibodies at 4°C. Dynabeads Protein G (Invitrogen) was used to precipitates antibody-bound proteins. Samples were separated by SDS-PAGE and electro-transferred to a polyvinylidene difluoride membranes (PVDF; Millipore). The membrane was blocked at room temperature for 1h and incubated with the indicated primary antibodies overnight at 4°C. The primary antibodies used were as follows: phospho-Serine (P5747, Sigma-Aldrich; 1:1000), phospho-Threonine (#9386, Cell signaling; 1:1000), DLX3 (PA5-40506, Invitrogen, OR, USA), HA(F-7, Santa Cruz Biotechnology), Flag (F3165; Sigma-Aldrich, USA; 1:5000) and α-tubulin (T5168, Sigma-Aldrich; 1:3000). Horseradish peroxidase-conjugated antibodies (Millipore) were used as secondary antibodies. The peroxidase reaction products were visualized with WESTZOL (Intron). All signals were detected by Amersham Imager 600 (GE Healthcare Life Sciences).

### Site-directed mutagenesis

Using the Flag-tagged wild-type *Dlx3* plasmid (Flag-DLX3 or DLX3^WT^) as a template, PCR-based mutagenesis was performed using a primer containing the desired mutation site (T134, S136, and S137). We generated mutations that mimicked either inactive phosphorylation status (serine/threonine to alanine) or activated phosphorylation status (serine/threonine to aspartic acid). The completed PCR product was cut with DpnI for 2 h and transformed into E.coli DH5alpha competent cells by heat shock. Colonies from each plate were grown and DNA was extracted. Mutations were verified by sequencing.

### Real-Time quantitative PCR (RT-qPCR)

For the RT-qPCR, the total RNA of the cells was extracted using TRIzol reagent (Invitrogen, Carlsbad, CA). The extracts were reverse transcribed using Maxime RT PreMix (#25081, iNtRON, Korea). RT-qPCR was performed using a StepOnePlus Real-Time PCR System (Applied BioSystems, USA). The amplification program consisted of 40 cycles of denaturation at 95°C for 15 s and annealing at 60 °C for 30 s. The expression levels of each gene are expressed as normalized ratios against the *B2m* housekeeping gene. RT-qPCR experiments were processed at least three independent trials with consistent results. The oligonucleotide primers for RT-qPCR are described in Supplementary Table 1.

### ChIP assay

Cells were cross-linked with 4% paraformaldehyde for 10 minutes at room temperature. Glycine was added to a final concentration of 125 mM for 5 minutes to quench the formaldehyde crosslinks. Cells were washed with ice-cold phosphate buffered saline, harvested by scraping, pelleted, and resuspended in SDS lysis buffer (50 mM Tris-HCl [pH 8.1], 1% SDS, 10 mM EDTA) with complete protease inhibitor cocktail (Roche). Cell extracts were sonicated with a Bioruptor TOS-UCW-310-EX (output, 250W; 25 cycles of sonication with 30-seccond intervals; Cosmo Bio). Samples were centrifuged at 14,000 rpm at 4°C for 10 minutes, and the supernatants were diluted 10-fold in dilution buffer (20 mM Tris-HCl [pH 8.0], 2 mM EDTA, 1% Triton X-100, 150 mM NaCl, and complete protease inhibitor cocktail). Chromatin samples were precleared with protein A-agarose beads (Santa Cruz) for 2h before immunoprecipitation against Flag (Sigma-Aldrich) antibodies overnight at 4 °C. Immune complexes were collected with protein A-agarose beads. Samples were washed five times (first wash with low salt immune complex wash buffer [20 mM Tris-HCl, pH.8.0, 2 mM EDTA, 1% Triton X-100, 0.1 % SDS, and 150 mM NaCl], second wash with high salt immune complex wash buffer [20 mM Tris-HCl, pH.8.0, 2 mM EDTA, 1% Triton X-100, 0.1% SDS, and 500 mM NaCl], third wash with LiCl immune complex wash buffer [10 mM Tris-HCl, pH.8.0, 1 mM EDTA, 250 mM LiCl, 1% NP-40, and 1% Na-deoxycholate], and the last two washes with TE buffer). Immunoprecipitated samples were eluted with buffer containing 1% SDS and 100 mM NaHCO3 at room temperature. Eluates were heated overnight at 65°C to reverse crosslinks after adding NaCl to a final concentration of 100 mM. Genomic DNA was extracted with a PCR purification kit (GeneAll). Precipitated chromatin by real-time PCR and the readouts were normalized using 5% input chromatin for each sample. The experiments were repeated two or more times. Primers were listed in Supplementary Table 2.

### Luciferase assay

The mHat9d cells were transiently transfected with 3X DRE-Luc and HA-MAST4 PDZ, 3Flag-DLX3 and series of mutation constructs by Neon Electroporation Transfection system (ThermoFisher). After 24 h, cells were lysed and the luciferase activities were analyzed using the Luciferase Assay System kit (Promega) according to the manufacturer’s protocol. All assays were done in triplicate, and all values were normalized for transfection efficiency against β-galactosidase activities.

### Statistics

Data analyses were performed in GraphPad Prism 8.0.2. All statistical analyses used nonparametric methods, which do not assume an underlying normal distribution in the data. Unpaired *t*-test was used for comparison of means from two experimental groups. All numerical bar graphs are represented as mean±SD. Throughout the paper, the *p*-values for the comparisons are given in the figure legends and denoted on the graphs according to the following key: \**p*≤0.05, \*\**p*≤0.01, \*\*\**p*≤0.001, \*\*\*\**p*≤0.0001, non-significant (ns) *p*>0.05.

## Data Availability

The RNA-Seq raw datasets are generated as FASTQ format. Total 4 raw data can be downloaded from the Sequence Read Archive (SRA) under BioProject accession number PRJNA785577. Reads alignment metadata and processed normalized gene expression data of RNA-Seq have been deposited in Dryad; https://doi.org/10.5061/dryad.4b8gthtf1.

## Acknowledgement

We are grateful to Mr. S. Kenmotsu and Mr. H. Sano for their technical assistance with EPMA and micro computed tomography analyses. This work was supported by the National Research Foundation of Korea (NRF) Grant funded by the Korean Government (MSIP)(NRF-2019R1A2C3005294). This research was supported by the Bio-&Medical Technology Development Program of the National Research Foundation (NRF) & funded by the Korean Government (MSIP&MOHW) (No. 2017M3A9E4048172).

## Figure Legends

**Figure 1 - Figure supplement 1.**
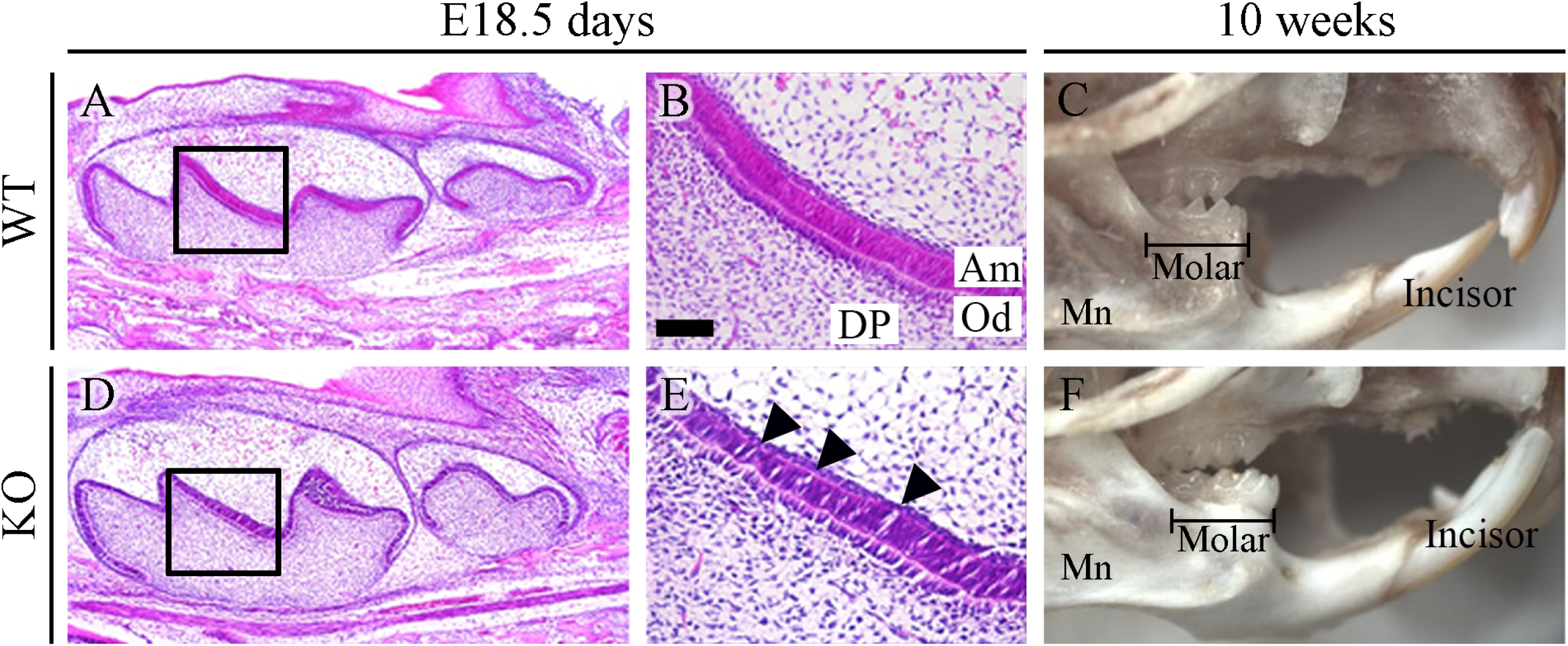
Molar development is not affected by the ablation of *Mast4*. (A, B, D, E) Mandibular molar tooth germ of E18.5 WT and *Mast4* KO mice. Bell-stage tooth germs have elongated and well-aligned ameloblast layers. (E arrowheads) Several defects in ameloblast alignment of *Mast4* KO tooth germ. (C, F) Mandibular bone and teeth of 10 weeks WT and *Mast4* KO mice after soft tissue removal. Contrary to the difference in length and direction of incisors, no difference was found between the WT and *Mast4* KO molars. Am; ameloblast, DP; dental papilla, Od; odontoblast, Mn; mandible, scale bars; B, E, 200 μm.

**Figure 1 - Figure supplement 2.**
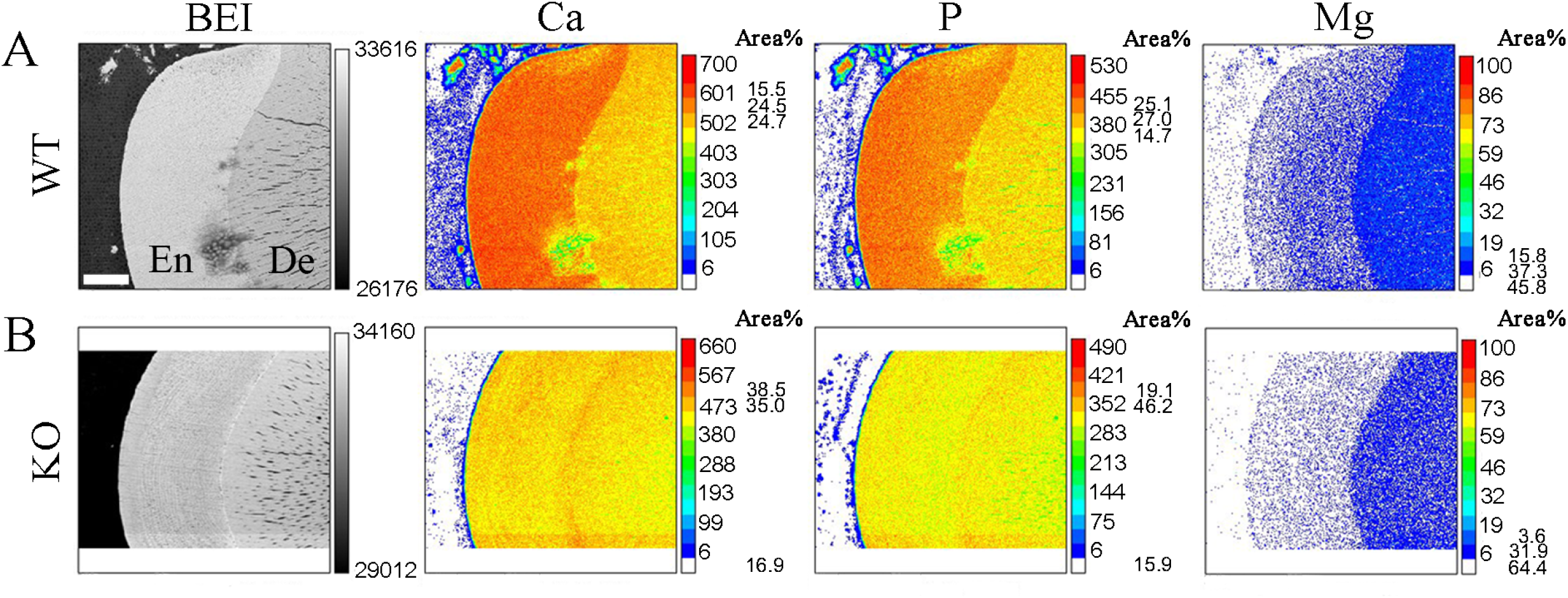
Mineral density of mandibular incisors. (A, B) Electron probe microanalyzer (EPMA) analysis of 6-week mandibular incisors. The mineral composition of the enamel is decreased in the *Mast4* KO incisors. Numbers in addition to the indicator bars are mineral intensity (a.u.). The area % values show the three most measured intensity ranges. BEI; backscattered electron image, Ca; calcium, P; phosphorus, Mg; magnesium. Scale bars; A, B, 50 μm.

**Figure 2 - Figure supplement 1.**
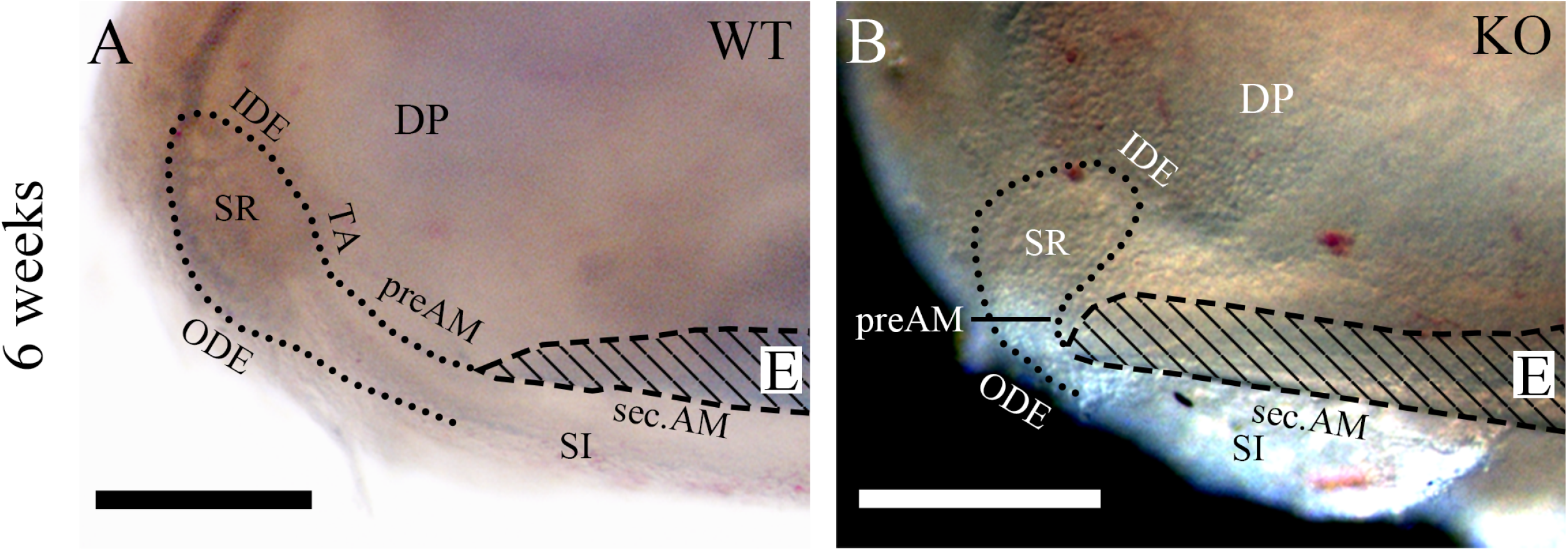
Labial cervical loops of mandibular incisors are exposed. Under a stereoscopic dissecting microscope, mandibles of 6-week WT and *Mast4* KO mice are dissected to expose the apical buds. (A) In the WT mandibular incisor, initiation of the enamel matrix was observed after the TA zone. (B) However, the initiation of enamel in the *Mast4* KO incisor was shifted to the apical side, and the cervical loop was reduced compared to the WT. Dotted lines; margin of apical buds, E; enamel, DP; dental papilla, sec.Am; secretory Ameloblast, preAm; Preameloblast, IDE; inner dental epithelium, ODE; outer dental epithelium, SR; stellate reticulum, SI; stratum intermedium, TA; transit-amplifying zone. Scale bars; A, B, 250 μm.

